# Armored Bicistronic CAR T Cells with Dominant-negative TGF-β Receptor II to Overcome Resistance in Glioblastoma

**DOI:** 10.1101/2024.02.26.582107

**Authors:** Nannan Li, Jesse L. Rodriguez, Yibo Yin, Meghan T Logun, Logan Zhang, Shengkun Yu, Kelly A. Hicks, Vicky Zhang, Laura Zhang, Chuncheng Xie, Jiabin Wang, Joseph A Fraietta, Zev A. Binder, Zhiguo Lin, Donald M. O’Rourke

## Abstract

Chimeric antigen receptor (CAR) T cells have shown significant efficacy in hematological diseases. However, CAR T therapy has demonstrated limited efficacy in solid tumors, including glioblastoma (GBM). One of the most important reasons is the immunosuppressive tumor microenvironment (TME), which promotes tumor growth and suppresses immune cells to eliminate tumor cells. The human transforming growth factor-beta (TGF-β) plays a crucial role in forming the suppressive GBM TME and driving the suppression of the anti-GBM response. In order to mitigate TGF-β mediated suppressive activity, we combined a dominant-negative TGF-β receptor II (dnTGFβRII) with our previous bicistronic CART-EGFR-IL13Rα2 construct, currently being evaluated in a clinical trial, to generate CART-EGFR-IL13Rα2-dnTGFβRII, a tri-modular construct we are developing for clinical application. We hypothesized that this approach would more effectively subvert resistance mechanisms observed with GBM. Our data suggests that CART-EGFR-IL13Rα2-dnTGFβRII significantly augmented T cell proliferation and enhanced functional responses, particularly in a TGFβ-rich tumor environment. Additionally, *in vivo* studies validated the safety and efficacy of the dnTGFβRII cooperating with CARs in targeting and eradicating GBM in a NSG mouse model.

## INTRODUCTION

Glioblastoma (GBM) is the most malignant and common primary brain tumor in adults. GBM has an extremely invasive nature, leading to infiltration of surrounding normal brain tissue. As a result, even with gross total resection, achieving a surgical cure remains impossible.^1^ GBM is therefore associated with an almost 100% recurrence rate, despite current standard-of-care treatment, including surgical resection, concurrent radiation therapy, and maintenance chemotherapy (Temozolomide).^2,3^ The median overall survival (OS) ranges from 14.6 months to 20.5 months.^4–6^ Currently, there is no approved second-line therapy for GBM patients. Given the incurable nature of GBM, the field needs new therapeutic strategies. Immunotherapy, and more specifically, chimeric antigen receptor (CAR) T cell therapy have shown significant efficacy in hematological diseases, with a recent study confirming the durability of CAR T cells in patients for up to ten years.^7^ This has prompted research into the use of CAR T cells as a potential treatment for solid tumors. However, to date, this approach has demonstrated limited efficacy in solid tumors, including GBM, with few positive results observed in CAR T clinical trials.^8–12^

We have completed two phase I clinical trials using CAR T cells for the treatment of GBM, targeting the epidermal growth factor receptor variant III (EGFRvIII).^11,13^ These trials allowed for evaluation of the surrounding tumor microenvironment (TME), pre- and post-infusion. Compared to pre-infusion tumor specimens, post-CAR T–infusion tumor specimens had markedly increased expression of multiple immunosuppressive molecules, including the human transforming growth factor-beta (TGF-β). TGF-β is consistently highly expressed in GBM tumor cells and patient tumor tissues.^14,15^ The TGF-β pathway has been identified as a significant contributor to glioma development due to its influence on various processes such as tumor cell proliferation, invasion, angiogenesis, and maintenance of stem cell stemness.^16–18^ Additionally, TGF-β plays a crucial role in driving the immune-suppression of the anti-GBM responses within the TME.^19,20^ It regulates a large number of immune cell types and controls both adaptive and innate immunity, thus forming a negative immune regulatory network.^21^ Given its dual role in both promoting tumor growth and suppressing immune cells, multiple strategies are currently being explored to mitigate the effects of TGF-β. These include targeting the latency-associated peptide,^22^ leveraging the α_v_ integrin/TGF-β axis,^23^ inhibiting the TGF-β^24^ or kinase receptor type I using small molecule inhibitors,^25^ employing TGF-β switch receptor,^26–28^ and TGF-β CAR T cells.^29^ However, these methodologies also raise considerable safety issues, as it has been confirmed that first-generation TGF-βRI inhibitors can result in cardiac toxicity.^30^ This toxicity risk becomes particularly concerning when advancing from preclinical research into the realm of clinical trials.

In order to mitigate TGF-β activity and decrease the side effect risk, previous groups have developed a truncated TGF-β receptor II.^31^ The truncated receptor lacks the signaling domain, creating a dominant-negative variant of TGF-β receptor II (dnTGFβRII) that inhibits the suppressive signal transduction pathway in the engineered T cells. The strategy of dnTGFβRII combined with CAR T cells has proven highly effective against prostate solid cancer in patients.^32,33^ Cytotoxic T lymphocytes cells expressing dnTGFβRII led to four of seven Hodgkin lymphoma patients achieving clinical responses, two of which have had a duration of more than 4 years.^34,35^ Importantly, the extracellular component of the receptor is not absent;^36–38^ it serves as a sink for TGF-β, assisting both engineered T cells and bystander T cells. This approach is believed to be safer than blocking the TGF-β’s normal physiological function.^32,33,35,39^

Given the tissue and signaling pathway heterogeneity seen within GBM, TGF-β blocking monotherapy will likely not lead to clinically relevant impacts.^40–42^ This underscores the importance of combinatorial targeted therapy. Here we combined the previous developed dual targeting CART-EGFR-IL13Rα2 (Interleukin 13 receptor alpha 2) with the dnTGFβRII construct to evaluate its potency both *in vitro* and *in vivo*. We propose the use of a dual targeting CART-EGFR-IL13Rα2 in conjunction with a dnTGFβRII construct as an effective approach to subvert resistance mechanisms found in the treatment of GBM.

## RESULTS

### TGF-β1 is highly expressed in GBM

Analysis of RNA-seq data from gliomas from the genomic data commons data portal on The Cancer Genome Atlas (TCGA) revealed that TGF-β1 expression is significantly higher in the GBM cohort compared to other glioma histology (p<0.0001) (Figure 1A). Further evaluation in GBM cases alone demonstrated TGF-β1 expressing patients had significantly worse OS than low TGF-β1 expressing patients (10.6 months vs 15.6 months, respectively, p=0.001344) (Figure 1B). In order to assess model fitness, media was collected from tumor cell lines and tested by ELISA to quantify the latent TGF-β secretion, using the prostate-specific membrane antigen (PSMA) tumor cell line, PC3, as a positive control (Figure 1C).^32^ U87MG and D270MG had high levels of TGF-β secretion and were chosen for the subsequent experimentation. GBM xenograft tumors stained with anti-TGF-β1 antibody via immunohistochemistry displayed high TGF-β1 expression within tumor tissue areas but not in adjacent normal brain regions (Figure 1D). Tissue sections from the mice spleen and cerebral cortex were used as positive and negative controls, respectively. These findings indicated TGF-β1 is highly expressed in malignant gliomas, combined with a lack of expression on normal brain tissues. Taken together, TGF-β is commonly found in GBM cells and represents a viable target to reduce immunosuppression.

**Figure 1.**
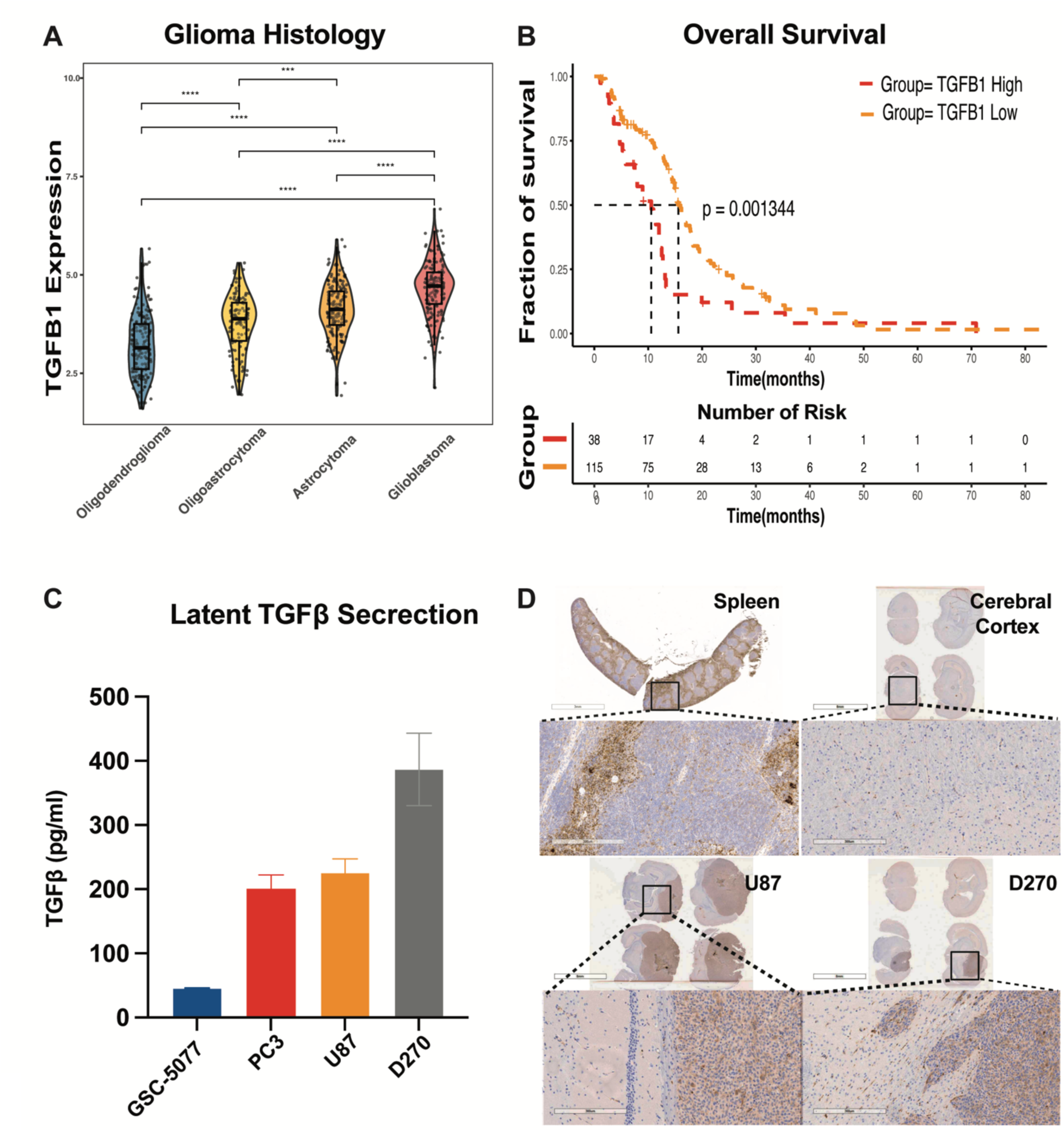
TGF-β1 is highly expressed in GBM. (A) TGF-**β**1 expression segregated by glioma histology based on RNA-seq data from the Cancer Genome Atlas (TCGA) database. The y-axis indicates the relative expression levels of each case. (B) Overall survival curve for high and low TGF-β1 expression group in accordance with the TCGA database, calculating by upper and lower quartile of the TGF-**β**1 expression in glioblastoma cases on Kaplan-Meier analysis. (C) Latent TGF-β secretion collected by glioblastoma cell lines supernants. (D) Immunohistochemical staining with anti-TGF-β1 antibody on tissue sections from NSG mice intracranially implanted with U87 and D270 gliomas. The spleen and cerebral cortex sections were used as positive and negative controls, respectively. Statistically significant differences were calculated by Kruskal-Wallis Test, ***p<0.001, ****p<0.0001.

### T cells expressing dnTGFβRII construct block immunosuppressive TGF-β signaling

To mitigate the immunosuppressive role of TGF-β1 in GBM, we combined a dominant-negative TGF-β receptor II molecule with our previous CART-EGFR-IL13Rα2 construct to generate CART-EGFR-IL13Rα2-dnTGFβRII, named as 806-Hu07-dnTGFβRII CAR, evaluating the potential of the dnTGFβRII molecule to bind TGF-β1 in the GBM TME. This binding would both abrogate the immunosuppressive TGF-β signal in transduced T cells and serve as a TGF-β1 sink, decreasing the amount of free suppressive factor in the tumor milieu available to bind to non-transduced T cells. We used 806-Hu07-mCherry CAR as a negative control (Figure 2A). For an additional irrelevant CAR control, we employed dnTGFβRII-M5, a clinical CAR T construct that targets mesothelin. This particular CAR was utilized to discern whether the T cells expressing truncated TGFβRII would induce toxicity to normal tissues.

**Figure 2.**
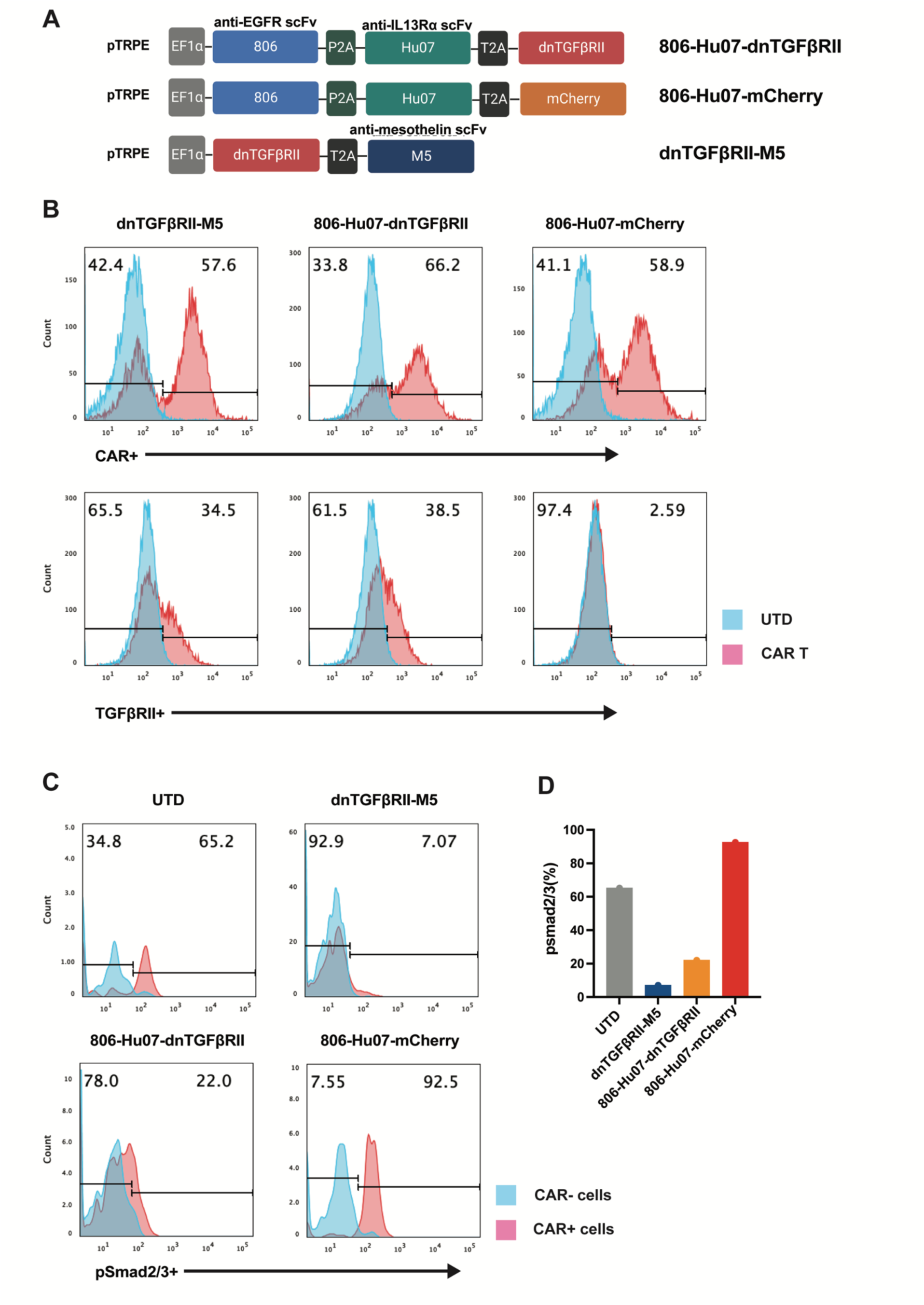
T cells expressing dnTGFβRII block immunosuppressive TGF-β signal. (A) Schematic vector maps of 806-Hu07-dnTGFβRII/806-Hu07-mCherry/dnTGFβRII-M5 CAR constructs. (B) Flow cytometric detection of T cell transduction. Cell populations were normalized to the same expression rate prior to experimentation. Upper row is stained for CAR expression by biotinylated protein L; lower row is stained for TGFβRII expression. (C) TGF-β signal induction of each group was assessed via intracellular phosphorylated Smad2/3 and quantified in (D).

Expression of the CAR was determined on human T cells transduced with lentiviral vectors, revealing approximately 60% CAR transduction among all CAR T cell groups expanded (Figure 2A). We also detected TGFβRII expression in CAR T cells, showing approximately 40% TGFβRII expression at the 806-Hu07-dnTGFβRII and dnTGFβRII-M5 groups (Figure 2B). All the T cells exhibited a similar fold expansion and resting volume kinetics during the manufacturing (Supplement 1A). To determine if the dnTGFβRII could block the immunosuppressive TGF-β/pSMAD signaling in T cells, we starved the T cells in culture for 24 hours and spiked active TGF-β1 into the culture media to induce TGF-β/pSMAD signaling. We found that both 806-Hu07-dnTGFβRII and dnTGFβRII-M5 CAR constructs blocked the formation of intracellular phosphorylated intracellular Smad2/3 (pSMAD) production, which is down-stream in the TGF-β signaling pathway (Figure 2C). The 806-Hu07-mCherry CAR and un-transduced T cells (UTD) groups readily generated intracellular pSMAD, confirming dominant negative-induced inhibition of TGF-β signaling (Figure 2C and D). These findings suggested that dnTGFβRII cell-surface expression on the 806-Hu07-dnTGFβRII CAR blocked the TGF-β immunosuppressive signaling pathway, did not interfere with CAR T cell expansion, and could potentially rescue T cells from the suppressive GBM microenvironment.

### dnTGFβRII boosts CAR T cells proliferation in the presence of TGF-β1 after long-term repeated stimulation *in vitro*

To evaluate the impact of dnTGFβRII in improving the activity of the bicistronic 806-Hu07-dnTGFβRII CAR construct, we co-cultured CAR T cells with D270MG GBM cells.^43^ However, no significant difference was observed in tumor cell killing between the 806-Hu07-dnTGFβRII and the control 806-Hu07-mCherry CAR cohorts (p=0.3686) (Supplement 1B). We hypothesized that 806-Hu07-dnTGFβRII CAR would perform better in an immunosuppressive media where TGF-β1 is present. To test this hypothesis, we conducted a cytotoxicity assay using either media alone or media conditioned with 20ng/ml of TGF-β1 media at the indicated effector: target (E: T) ratios (Figure 3A). However, we did not observe a significant difference in cytotoxic activity between the 806-Hu07-dnTGFβRII CAR and the control 806-Hu07-mCherry CAR group. Moreover, these results were also confirmed via detection of expression of T cell short-term activation markers CD69 and CD25 when co-cultured CAR T cells with U87MG-EGFRvIII (U87vIII) cells for 20 hours (Figure 3B, Supplement 1C).^44^

**Figure 3.**
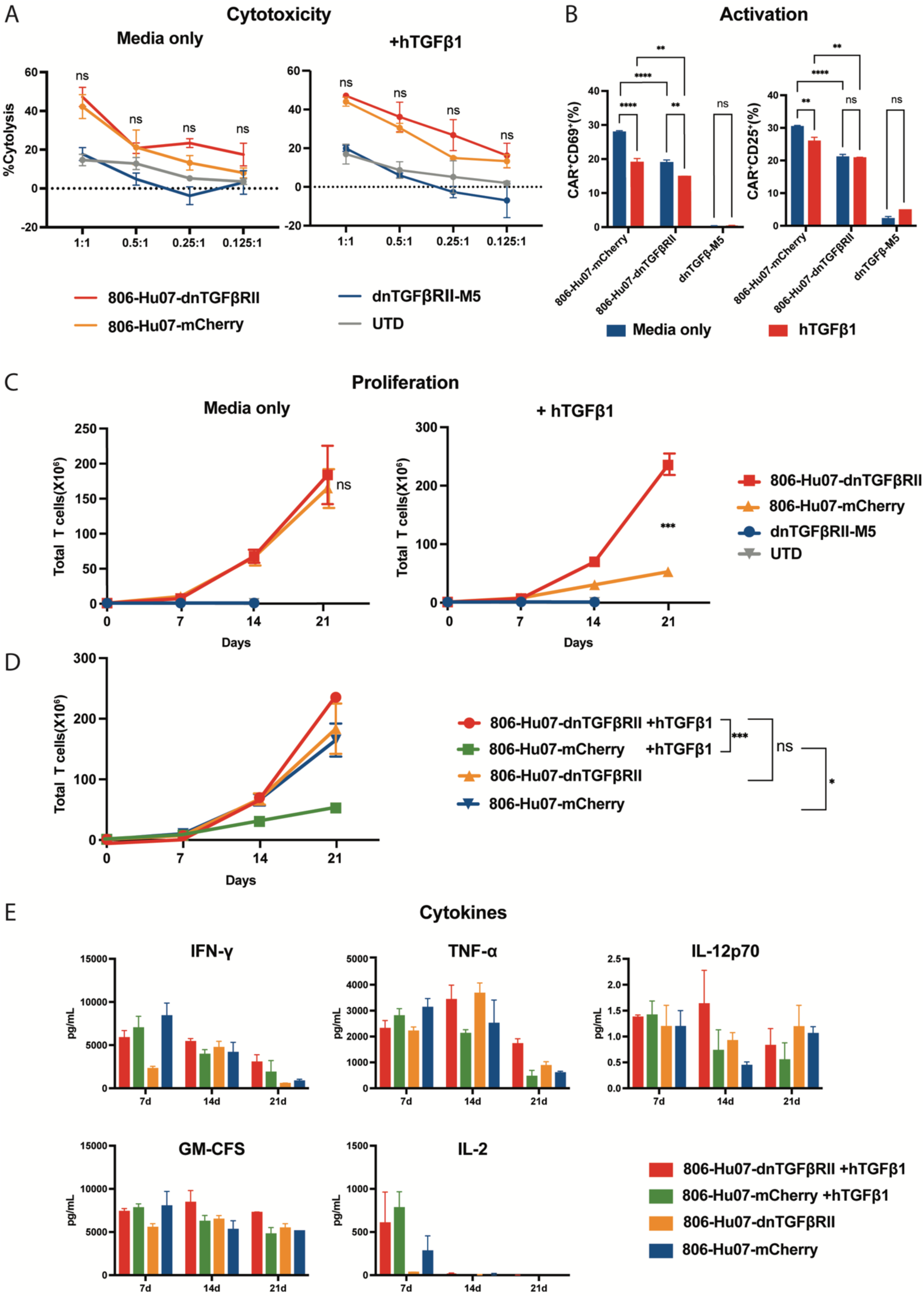
dnTGFβRII boosts CAR T cells proliferation in the presence of hTGF-β after long-term repeated stimulation *in vitro.* (A) In vitro cytotoxicity after 16 hrs of co-culture with the U87vIII-CBG-GFP tumor cells, demonstrating no significant difference along with either 20ng/ml of TGF-β1 or media only at indicated E:T ratios. Statistically significant differences were calculated by ordinary one-way ANOVA with Tukey’s multiple comparisons test. (B) Short-term activation quantified by CD69 and CD25 expression in CAR+ T cells after 20 hrs of co-culture with the U87vIII tumor cell line, which was conditioned with 20ng/ml of TGF-β1 or media only. Statistically significant differences were calculated by two-way ANOVA with Tukey’s multiple comparisons test. (C) CAR T cells were restimulated by irradiated U87vIII tumor cell lines with either conditioned media or media only every week as shown up. (D) The comparison of proliferation between 806-Hu07-dnTGFβRII and 806-Hu07-mCherry CAR T cells that were either in conditioned media or media only. (E) Collected the supernatant from proleferation coculture plates on days 7, 14, and 21, then stored it at -80 degrees celsius for future analysis of cytokine secretion. Proliferation assay was calculated by unpaired t test with two-tailed. ns, not significant; *p<0.05, **p<0.01, ***p<0.001, ****p<0.0001. Data are presented as means ± SEM.

We reasoned that short-term stimulation may not physiologically reflect the suppressive effects of TGF-β on CAR T cell function so we shifted to a long-term U87vIII co-culture model. In the presence of 20ng/ml of TGF-β1 conditioned media, the proliferation of 806-Hu07-mCherry CAR T cells was significantly suppressed (p=0.0007) when compared with 806-Hu07-dnTGFβRII CAR T cells (Figure 3C). This suggested that the dnTGFβRII construct helped overcome the immunosuppressive conditions and retain a proliferation capacity comparable to the media-only group (p=0.32) (Figure 3D). Conversely, the 806-Hu07-mCherry CAR T cells lacked this adaptability, displaying reduced proliferation in the tumor-like environment (p=0.0123).

In addition to improved proliferation, the 806-Hu07-dnTGFβRII CAR T cells also secreted a higher quantity of effector cytokines than the 806-Hu07-mCherry CAR T cells, including IFN-γ, TNF-α, IL-12, GM-CSF, and IL-2 (Figure 3E). This was observed in the conditioned media that was collected from the weekly stimulated proliferation assay at later stages, specifically at days 14 and 21. Notably 806-Hu07-mCherry CAR T cells demonstrated more effector cytokine secretion than the 806-Hu07-dnTGFβRII CAR at Day 7. These results suggested that the enhanced response of the 806-Hu07-dnTGFβRII CAR T cells was not reliant on initial activation, but rather on long-term proliferation and longer kinetics of cytokines activation. These features may be important for T cell immunotherapies to ensure the expansion and cytotoxicity action against solid tumors.

### 806-Hu07-dnTGFβRII changes to an effector cell phenotype

To better understand the factors influencing the enhanced proliferation capacity of 806-Hu07-dnTGFβRII CAR T cells, we conducted a phenotype change assessment using T cells before and after the long-term repeated stimulation (Figure 4A). For Day 0, all the cohorts had similar T cell phenotypes, with the exception of the 806-Hu07-mCherry CAR T cells, which had more CD4+ “central memory” T cells than 806-Hu07-dnTGFβRII CAR T cells (p=0.0002) (Supplement 2A). During the long-term repeated stimulation assay, there was not much difference between the 806-Hu07-dnTGFβRII and 806-Hu07-mCherry CAR T cells in the media-only group after Day 14 and Day 21 co-culture. However, when exposed to media containing exogenous TGF-β1, these CAR T cells exhibited divergent trends. The 806-Hu07-mCherry CAR T cells retained a higher proportion of “central memory” cells with some “naïve” cells. However, the 806-Hu07-mCherry CAR T cells showed less-than-expected expansion when compared to the 806-Hu07-dnTGFβRII CAR T cells, despite the higher proportion of “central memory” cells.

**Figure 4.**
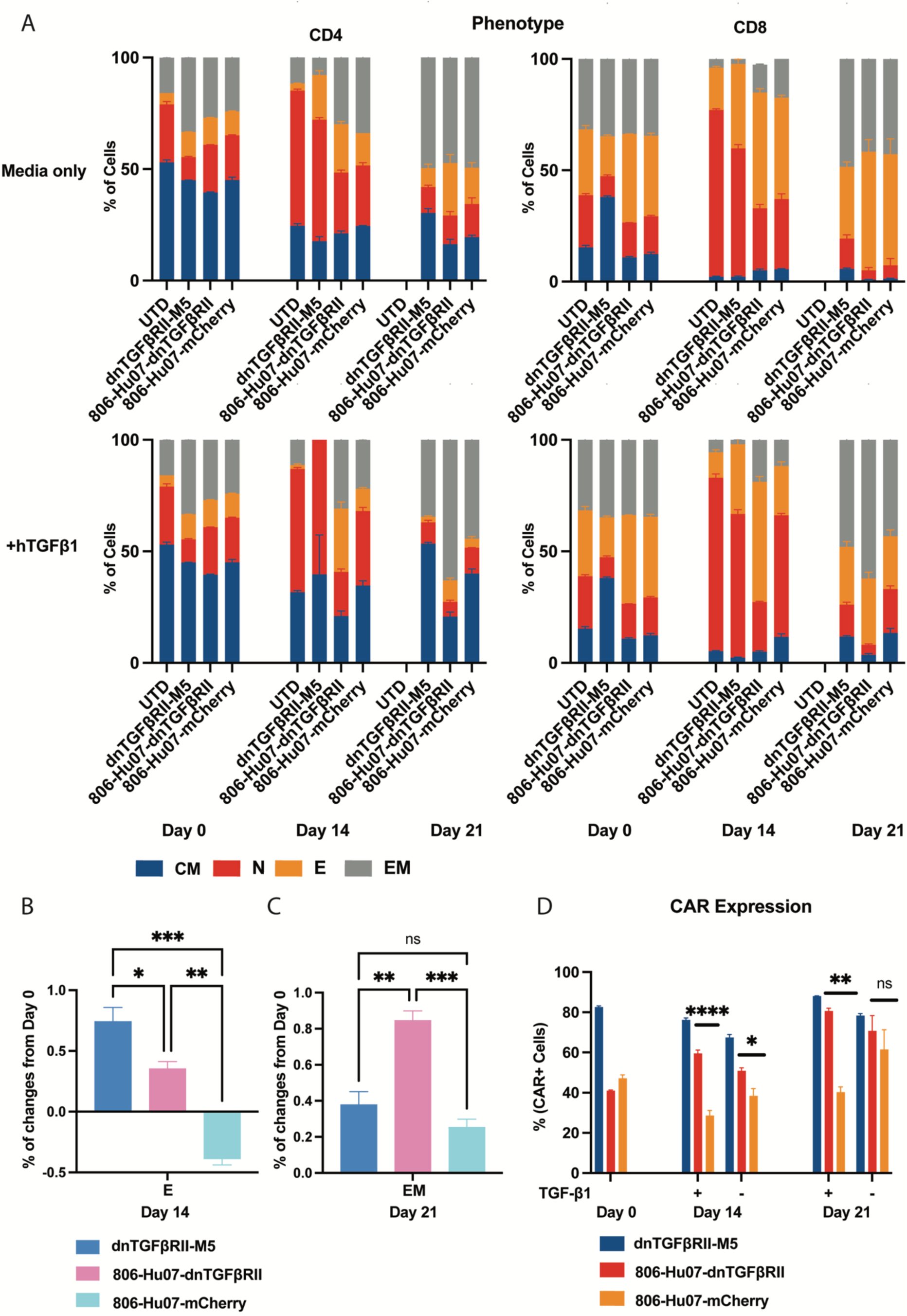
806-Hu07-dnTGFβRII changes to an effector cell phenotype. (A) *In vitro* long-term restimulation assay was performed weekly, with T cells being collected and stained to assess their phenotypic evolution over time, specifically on Day 0, Day 14, and Day 21. These changes were evaluated within different subsets: Central Memory (CM), Naive (N), Effector (E), and Effector Memory (EM) T cells. (B) The most notable alteration occurred in the cohort subjected to 20ng/ml of TGF-β1. On Day 14, the 806-Hu07-dnTGFβRII cells demonstrated a significantly more pronounced effector T cell phenotype than the 806-Hu07-mCherry CAR T cells (p=0.0015). (C) By Day 21, the final time point, the 806-Hu07-dnTGFβRII cells exhibited an enhanced “effector memory” phenotype (p=0.0009). (D) The CAR expression in both 806-Hu07-dnTGFβRII and 806-Hu07-mCherry CAR T cells was compared. These cells were subjected to either conditioned media (+) or media alone (-), and this comparison was conducted at the same time points as above. The CAR expression was nomilized to 41% at Day 0 to start later assays. Statistical significance was determined through a two-way ANOVA at (A) and (D), and a ordinary one-way ANOVA at (B) and (C), supplemented by Tukey’s multiple comparisons test. The following significance levels were used: ns denotes not significant, *p<0.05, **p<0.01, ***p<0.001, ****p<0.0001. Data are presented as the mean ± Standard Error of the Mean (SEM).

In contrast, the 806-Hu07-dnTGFβRII construct not only demonstrated increased proliferation in the tumor-simulating milieu but also exhibited a significant “effector” phenotype during repeated stimulation at the intermediate stage (p=0.0015) (Day 14), likely contributing to its tumor-killing potential (Figure 4B). By Day 21, “effector” transitioned into an “effector memory” phenotype, allowing the T cells to simultaneously retain cytotoxicity and memory ability (p=0.0009) (Figure 4C). Moreover, the CAR expression of these T cells confirmed that the 806-Hu07-dnTGFβRII construct maintained more stable CAR expression than the 806-Hu07-mCherry construct, particularly under immunosuppressive conditions (Day 14, p<0.0001; Day 21, p=0.016; Figure 4D). These characteristics likely explained how the 806-Hu07-dnTGFβRII CAR T cells to sustain their potent cytotoxicity at the intermediate stage and a trend of increased proliferation during the late stages of the co-culture process.

### 806-Hu07-dnTGFβRII CAR T cells is safety to be used *in vivo*

After confirming the *in vitro* responsiveness of the 806-Hu07-dnTGFβRII CAR T cells, we proceeded to assess their anti-tumor efficacy *in vivo*. A key concern was whether incorporating the dnTGFβRII construct into CAR T cells might trigger toxicity in normal tissues. To address this, we introduced the glioma cell line D270-CBG-GFP subcutaneously into a NOD SCID gamma (NSG) mouse model. Seven days post-implantation, we treated the mice with an intravenous (IV) infusion of CAR T cells (Figure 5A). We utilized bioluminescent imaging (BLI) every 3-4 days to track tumor growth. CAR T cell toxicity was assessed based on mice weight changes. Over the course of 47 days, the 806-Hu07-dnTGFβRII CAR T group displayed no difference in mice body weights compared to the control groups 806-Hu07-mCherry CAR, dnTGFβRII-M5, or UTD (Figure 5B). Additionally, there were no reported incidents of hair loss or ruffling, rash, or reduced mobility. These findings suggested that the dnTGFβRII construct did not precipitate graft-versus-host disease (GvHD). Furthermore, the dnTGFβRII construct did not compromise the cytotoxic capabilities of CAR-806-Hu07, as it demonstrated a comparable anti-tumor response in this mouse model (p>0.9999) (Figure 5C). BLI data highlighted the potency of both the 806-Hu07-dnTGFβRII and 806-Hu07-mCherry CARs in targeting subcutaneously injected glioma cells (Figure 5D). In contrast, the UTD control was ineffective and the dnTGFβRII-M5 CAR did not manifest non-specific cytotoxic effects. In conclusion, the 806-Hu07-dnTGFβRII CAR was safe for *in vivo* use and CAR T cells expressing this tri-modular construct maintained the robust cytotoxic capabilities.

**Figure 5.**
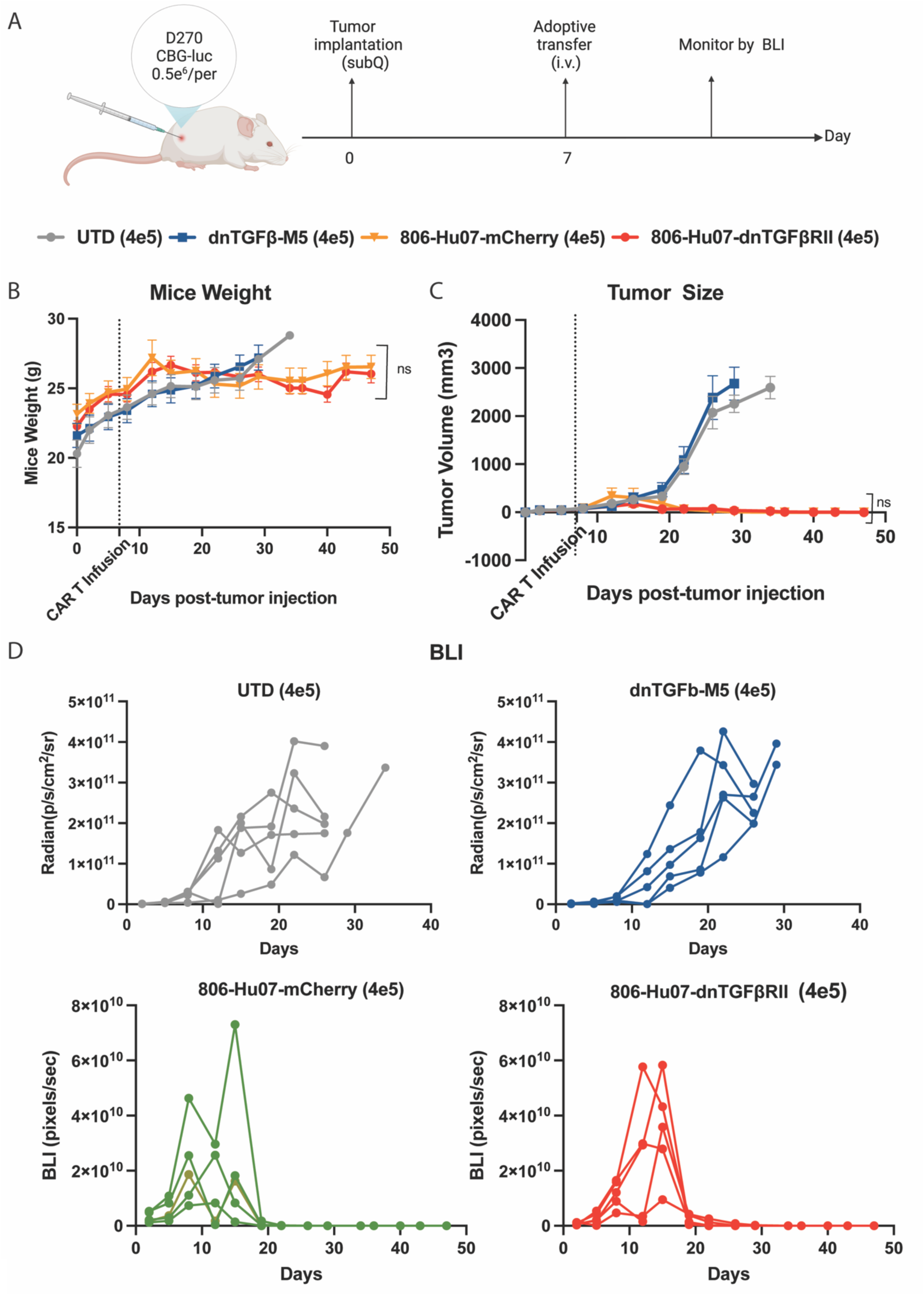
dnTGFβRII construct is safety *in vivo.* (A) Schematic of the D270-CBG-GFP NSG mice model (n = 5 per cohort). 5e5 tumor cells were implanted subcutaneously and treated by intravenous infusion of CAR T cells into each mouse, 7 days after tumor implantation. The BLI of tumor burden was performed 3-4 days. (B) Mice weight were measured every 3-4 days. (C) Tumor size was measured by the calipers in length and width as the area of tumor. (D) Tumor regression was compared between each treated mouse by using BLI. Statistically significant differences at each time point were calculated by ordinary one-way ANOVA with Tukey test. ns, not significant. Data are presented as means ± SEM.

### 806-Hu07-dnTGFβRII CAR T cells enhance eradication GBM tumors *in vivo*

Having established the safety profile of the dnTGFβRII construct, we proceeded to test it in a more aggressive intracranial GBM mouse model, simulating the complex tumor microenvironment in which TGF-β is secreted by both tumor cells and the surrounding stroma.^45,46^ This was to determine whether the 806-Hu07-dnTGFβRII CAR T cells offered a more potent therapeutic advantage over the 806-Hu07-mCherry CAR T cells *in vivo*. We intracranially implanted U87vIII glioma cells, modified to express both EGFRvIII and the reporter gene CBG-GFP, into NSG mice. Eight days post-implantation, these mice were administered an IV infusion of the respective CAR T cells (Figure 6A).

**Figure 6.**
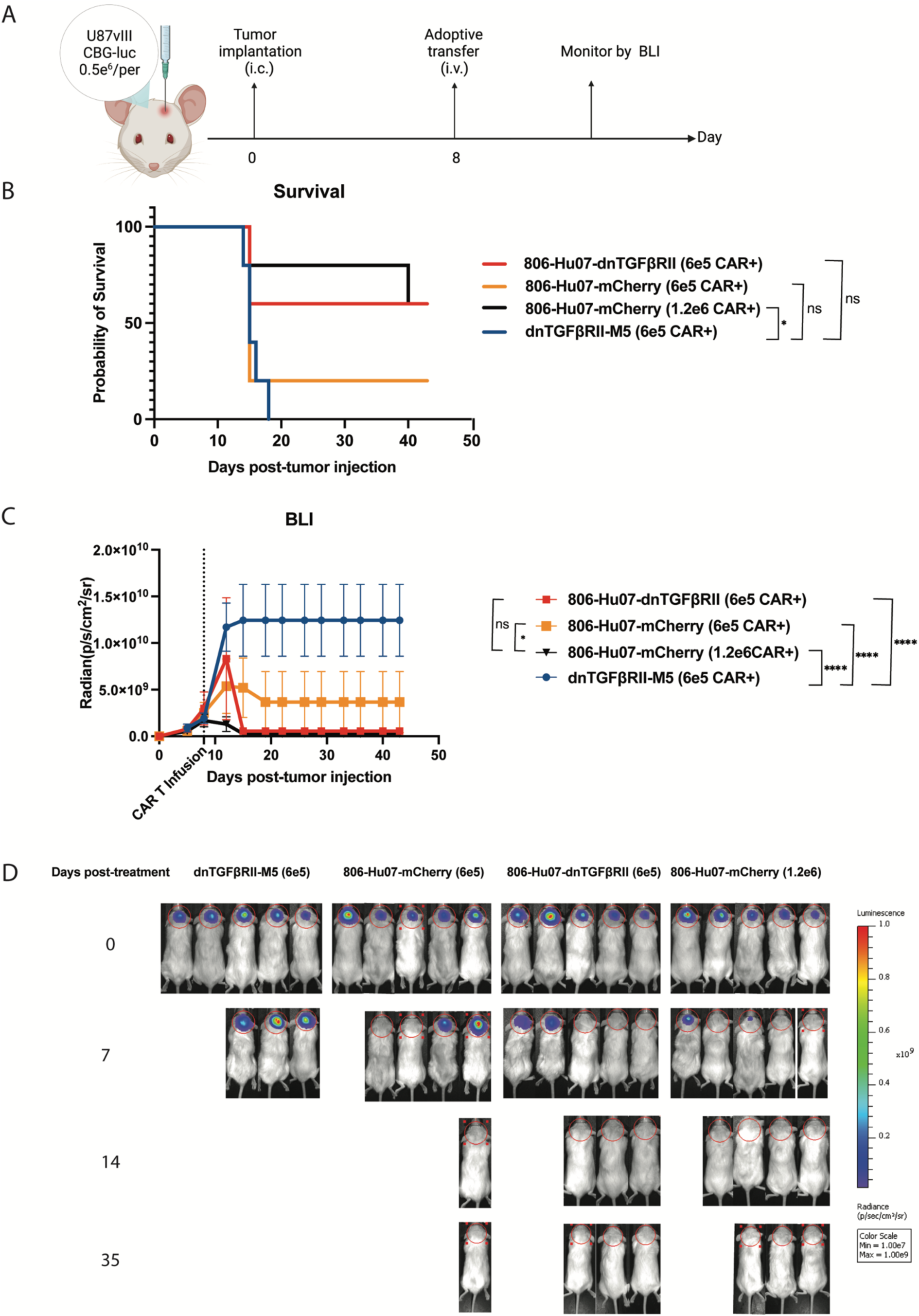
806-Hu07-dnTGFβRII CAR T cells eradicate GBM tumors *in vivo.* (A) Schematic of the U87vIII CBG-GFP NSG mice model (n = 5 per cohort). 5e5 tumor cells were implanted intracranially and treated by intravenous infusion of CAR T cells into each mouse, 8 days after tumor implantation. The BLI of tumor burden was performed 3-4 days. (B) Survival curves of treated mice were computed using the log-rank test with P values adjusted by the Bonferroni correction for multiple comparisons survival, based on time to experimental endpoint, was plotted using a Kaplan-Meier curve. (C and D) Tumor size was compared between each treated mouse by using BLI. Statistically significant differences of BLI at each time point were calculated by ordinary one-way ANOVA with Tukey test. Statistically significant differences of survival curves were determined using log-rank test. ns, not significant; *p<0.05, ****p<0.0001. Data are presented as means ± SEM.

To determine if dominant negative TGF-β receptor II generated resistance to the immunosuppressive TME onto CAR T cells *in vivo*, we used a sub-therapeutic dose of 0.6e6 CAR+ cells to treat each tumor-bearing mouse. Our previous study demonstrated that a dose of 1.2e6 806-Hu07-mCherry CAR+ cells can eradicate GBM orthotopic tumors.^43^ As we assumed, only the positive control group can promote the mice survival (p=0.0157) (Figure 6B). Surprisingly, mice treated with a sub-therapeutic dose of 806-Hu07-dnTGFβRII CAR T cells exhibited equivalent tumor eradication ability to the full-dose 806-Hu07-mCherry CAR group (p=0.8082), but 806-Hu07-mCherry with the sub-therapeutic dose failed (p=0.0252) (Figure 6C). Moreover, both the 806-Hu07-dnTGFβRII CAR and the full-dose positive control group eliminated tumor burden when compared with the negative control dnTGFβRII-M5 CAR (Figure 6D). In conclusion, dnTGFβRII not only rendered 806-Hu07-dnTGFβRII CAR T cells resistant to the immunosuppressive TME, but also amplified its proliferation ability and anti-tumor efficacy in *in vivo* orthotopic experiments.

## DISCUSSION

In order to counteract antigen escape and mitigate TGF-β signaling within the challenging GBM TME, we have developed CART-EGFR-IL13Rα2-dnTGFβRII T cells, also known as the 806-Hu07-dnTGFβRII CAR T cells. Our research indicates that these engineered CAR T cells dramatically outperforms the TGF-β-sensitive CAR (806-Hu07-mCherry) in terms of T cell proliferation *in vitro* and anti-tumor function *in vivo*. Additionally, dnTGFβRII augments T cell fitness by maintaining CAR expression, fortifying the effector functionality, and promoting effector cytokine production *in vitro* when exposure to the exogenous TGF-β1. Remarkably, 806-Hu07-dnTGFβRII CAR is safe to be used and displays efficacy in clearing advanced gliomas in an *in vivo* suppressive milieu.

It is well-established that the proliferative response of CAR T cells is a key predictor of clinical efficacy. However, in solid tumor trials, CAR T cell expansion is often limited.^47,48^ Our data reveals that the 806-Hu07-dnTGFβRII CAR T cells exhibit significant proliferative capabilities in both the media-only or exogenous TGF-β1 conditions during extended *in vitro* long-term repeated stimulation studies (Figure 3C). These data suggests that this construct can facilitate resistance to the anti-proliferative effects of exogenous TGF-β1. In contrast, the parental CAR T cells, 806-Hu07-mCherry, which retains TGF-β/SMAD signaling (Figure 2C) failed in the presence of TGF-β conditioned media (Figure 3D). The blockade of TGF-β has been confirmed to boost immune cells proliferation in GBM tumor models,^49,50^ underscoring the significance of a truncated TGF-β receptor II to render CAR T cells insensitive to the immunosuppressive agent TGF-β. Expressing dnTGFβRII not only shields transduced T cells but also benefits non-transduced cells.^35^ dnTGFβRII acts as a sink, effectively sequestering TGF-β from the tumor TME, thereby amplifying the protective effect for surrounding immune cells.

To further interrogate the enhanced proliferation mechanisms when blocking CAR T cells’ TGF-β signal, we sampled the co-culture media from long-term repeated stimulation assay. Analysis of this media revealed that 806-Hu07-dnTGFβRII CAR T cells secreted elevated levels of effector cytokines, including IFN-γ, TNF-α, IL-12, and GM-CSF, as evidenced in Figure 3E. However, the 806-Hu07-mCherry CAR T cells had decreased secretion both in the presence or absence of TGF-β conditioned media at the day 14 and day 21. These results are consistent with the demonstrated study that TGF-β represses T cells to secret the cytokines,^49,51^ the 806-Hu07-dnTGFβRII CAR T cells could block the TGF-β suppression. Given that 806-Hu07-dnTGFβRII and 806-Hu07-mCherry share the same antigen recognition domains, there should be no discernible difference in their activation profiles,^26^ as illustrated in Figure 3B. These results suggested that the enhanced response of the 806-Hu07-dnTGFβRII CAR T cells may be not reliant on initial activation. So, introducing the dnTGFβ receptor II would create a distinct proliferation and advantage with cytokine activation kinetics in the later stages of tumor elimination.

It is well known that TGF-β plays an important role in the process of tumor immune evasion by inhibiting the function of cytotoxic T cells.^52,53^ But the 806-Hu07-dnTGFβRII group exhibited a robust “effector” phenotype (Figure 4B), even under conditions mimicking a TGF-β-enriched tumor environment. This increased “effector” T cell phenotype helped 806-Hu07-dnTGFβRII CAR T cells eradicate tumor cells more rapidly, which was observed when we co-cultured CAR T cells and GBM tumor cells. What is particularly intriguing is that by Day 21, the T cell phenotype had transitioned from an “effector” to an enriched “effector memory” phenotype. The “effector” to an enriched “effector memory” phenotype endowed T cells with sustained effector functionality combined with improved T cell proliferation. Delving deeper into the mechanism, we observed that these 806-Hu07-dnTGFβRII CAR T cells displayed increased CAR

T expression (Figure 4D). Notably, even when exposed to exogenous TGF-β, there was no downregulation of CAR expression on these CAR T cells. This suggests that the genetically engineered CAR remains impervious to downregulation mediated by TGF-β.

There are also proactive approaches to target GBM by translating TGF-β immunosuppressive signal to the stimulant signal, such as TGF-β CAR.^29,49^ We propose that dnTGFβRII construct provides a relatively safe approach to counteract the GBM TME. Our findings were reassuring in that the dnTGFβRII construct safety was shown by the stability of mouse weights at the 806-Hu07-dnTGFβRII group, suggesting a lack of GvHD (Figure 5B). Furthermore, unspecific dnTGFβRII-M5 CAR showed no nonspecific toxicity activity, highlighting the safety of dnTGFβRII construct itself and without lymphoproliferative disorders in mouse models.^54,55^ 806-Hu07-dnTGFβRII and 806-Hu07-mCherry T cells had equivalent potent antitumor responses, these results suggested that the 806-Hu07-dnTGFβRII structure is effective and safe in treating GBMs. The 806-Hu07-dnTGFβRII construct was also resistant to the GBM TME and enhanced the antitumor response *in vivo* in intracranial mouse model. The dnTGFβRII construct improves cytotoxicity function of T cells also validated when paired with an anti-EGFRvIII CAR in glioma mouse models,^51^ as well as in clinical trials in other tumor models.^33,35^ In conclusion, the 806-Hu07-dnTGFβRII CAR T cells not only render the robust 806-Hu07-mCherry CAR resistant to the GBM TME but also amplify its and anti-tumor efficacy in *in vivo* experiments.

In conclusion, we synergized dnTGFβRII with our bicistronic CART-EGFR-IL13Rα2, which showed a remarkable ability to eliminate advanced glioma tumors in murine models and the Phase I clinical trial, ID NCT05168423, is currently ongoing at the hospital of University of Pennsylvania. Our research highlighted the tri-modular CART-EGFR-IL13Rα2-dnTGFβRII ability to significantly augment T cell proliferation and enhanced their functional response, especially in a TGFβ-rich tumor environment. Additionally, our *in vivo* studies validated the safety and efficacy of the dnTGFβRII in targeting and eradicating advanced gliomas in the mouse models.

## MATERIALS AND METHODS

### Cell Lines and Culture

The human GSC line (5077) was derived from excised tumor tissue obtained from the University of Pennsylvania Institutional Review Board with written informed consent from the patients (Department of Neurosurgery, Perelman School of Medicine, Philadelphia, PA). It was maintained in DMEM F12 Ham supplemented with penicillin/streptomycin, GlutaMAX-1, B27 minus A, epidermal growth factor, and basic fibroblast growth factor (Corning, Corning, NY). The U87MG cell line was obtained from the American Type Culture Collection (ATCC HTB-14) and cultured in MEM containing GlutaMAX-1, HEPES, pyruvate, penicillin/streptomycin (Thermo Fisher Scientific, Carlsbad, CA), and supplemented with 10% fetal bovine serum (FBS). This cell line was engineered to express the EGFRvIII protein, click beetle green luciferase, and green fluorescent protein through single-cell purification and expansion. The PC3 prostate cancer cell line was obtained from the ATCC (CRL-1435) and cultured in D10 media consisting of DMEM supplemented with 10% FBS, HEPES, penicillin, and streptomycin. The D270 glioma cells were grown and passaged in the flanks of NSG mice. The human glioma cell line U251 was provided by Dr. Jay Dorsey (Department of Radiation Oncology, University of Pennsylvania). The cells were routinely screened for identity and mycoplasma contamination.

### Immunohistochemical Stain

Immunohistochemical staining used a recombinant anti-TGFβ1 antibody (ab215715, Abcam, Waltham, Boston) and DAPI on tissue sections derived from NSG mice. These mice had been intracranially implanted with U87 and D270 gliomas, a procedure facilitated by Daniel Martinez (Pathology Core Laboratory of the Children’s Hospital of Philadelphia Research Institute, PA). Tissue sections of the spleen and cerebral cortex of mice were utilized as positive and negative controls, respectively.

### Vector Design

The 806-Hu07-mCherry CAR was assembled by combining an EGFR-targeting scFv (806) and an IL13Rα2-targeting scFv (Hu07), which were synthesized and ligated into a pTRPE lentiviral vector with the EF1alpha promoter. The construct ended with an mCherry (Twist Bioscience, San Francisco, CA). The dnTGFβRII construct was digested using AvrII and SalI enzymes and ligated into the 806-Hu07-mCherry construct at the same enzyme sites, replacing the mCherry gene to create the 806-Hu07-dnTGFβRII CAR construct. The CAR-dnTGFβRII-M5 was a gift from Joseph A. Fraietta at the Perelman School of Medicine, University of Pennsylvania. The control CAR-CD19 was a gift from Carl H. June lab, University of Pennsylvania. All CAR molecules have the same second-generation CAR design containing 4-1BB co-stimulatory domain with CD3 zeta chain.

### Lentiviral Vector Production and T Cell Transduction

Lentiviruses were packaged in HEK 293 T cells using a split genome approach and tittered using SUPT1 cells from ATCC (CRL-1942). Normal human T cells were isolated from the PBMC of the Human Immunology Core at the University of Pennsylvania and were transduced using lentiviral vectors. T cells were stimulated for 5 days with Dynabeads Human T-Activator CD3/CD28 (Life Technologies, Carlsbad, CA) at a bead-to-cell ratio of 3:1. The cell concentration was determined using a Coulter Multisizer (Beckman Coulter, Brea, CA) and maintained at 0.7e6 cells per mL until fully rested at approximately 300fl in volume. The cells were cultured in R10 media (RPMI-1640 supplemented with GlutaMAX-1, HEPES, pyruvate, penicillin/streptomycin, and 10% FBS) with 30 IU/mL rhIL-2 (Thermo Fisher Scientific, Carlsbad, CA). The CAR T cells were then cryopreserved in a mixture of 90% FBS and 10% DMSO for future use.

### TGF-β ELISA

The tumor cell lines were cultured at a density of 3e6 cells per flask for 3 days. The conditioned medium was collected by centrifugation, and stored at -80°C. The culture medium containing 10% fetal bovine serum is expected to have extra TGF-β1 secretion. Thus, a control medium should be used as a blank and subtracted from the samples. The thawed conditioned medium was processed using the TGF-β1 ELISA kit (DY240-05, R&D, Minneapolis, MN), and the samples were diluted four-fold prior to activation of latent TGF-β1 into an immunoreactive form.

### Bioluminescence Cytotoxicity Assay

The U87vIII-CBG luciferase target cells were co-cultured with effector T cells at different effector: target ratios for 16 hours, with either 20ng/ml of hTGF-β1 or media only. Prior to measuring the luminescence, 15μg of D-Luciferin (Gold Biotechnology, St. Louis, MO) was added and incubated at room temperature for 10mins. The luminescence was measured using a BioTek Synergy H4 hybrid multi-mode microplate reader. When calculating the cytotoxicity results, it was established that the target cells alone exhibited 0% lysis, the maximum lysis was observed in the treatment with 5% SDS Solution (Thermo Fisher Scientific, Carlsbad, CA).

### Proliferation Assay

The U87vIII cells were subjected to 10,000 rad of irradiation for 40min. Subsequently, triplicated cocultured of 0.2e6 tumor cells and 1e6 T cells in 12-well plates containing 4 ml of R10 media, with the addition of either 20 ng/ml of human TGF-β1 or media alone. The media was replenished if it became yellowed around day 4 or 5. Supernatant of 1-2 ml was collected from each well at day 7, 14, 21 centrifuged and stored at -80°C for subsequent cytokine analysis (Eve technologies, Calgary, Canada). The T cells were then resuspended, counted using a Beckman Coulter Multisizer 3 Cell Counter, and 1e6 T cells were transferred to the new 0.2e6 irradiated U87vIII cells to continue performing the proliferation assay weekly, the rest T cells were stained for phenotype and CAR expression.

### Impedance Cytotoxicity Assays

2e5 target tumor cells were seeded into Axion Biosystems microelectrode-containing 96-well plates (Axion Biosystems, Atlanta, GA). Each impedance plate was prepared prior to experimentation by coating with 20 μg/mL laminin overnight at 37°C. After coating was complete, wells were rinsed with diH2O and then overlaid with 100 μL of cell culture media. The plate was placed into the Axion Biosystems ZHT analyzer (Axion Biosystems, Atlanta, GA) to record baseline readings of the background impedance without cells present. After baseline was established, the plate was removed from the analyzer and was seeded with 5e5 target cells in a volume of 200 μL/well. After cell plating, the plate was left in the cell culture hood for 1h at room temperature to ensure settling and attachment of the cells down to the microelectrodes on the bottom surface. The plate was then returned to the analyzer and data collection began. Data were collected every 1 min for 24 h for cell monolayer growth measurement. For cytotoxicity assessment, the instrument was paused at 24h, and media was exchanged for media containing 1:1 dosages of effector cells or UTD control T cells, or media alone. Changes in impedance are reported as the resistive component of the complex impedance, as described previously. Using AxIS Z software (Axion Biosystems, Atlanta, GA), all data are corrected for “media alone” to remove small changes in media only impedance over time and then normalized to the impedance at the time of addition of effector cells. The % cytolysis calculations utilize the no treatment control and full lysis controls to determine % of target cell cytolysis as follows:

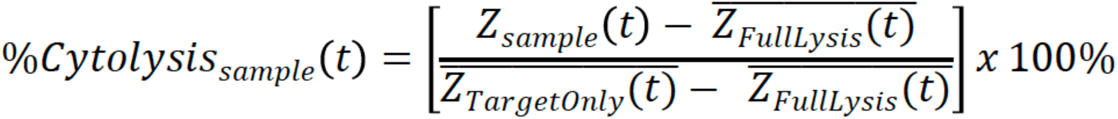

### Flow Cytometry

A 5-laser LSRFortessa flow cytometer was used for analysis. The cells were first stained with a live/dead viability stain (Thermo Fisher Scientific, Carlsbad, CA) in PBS. Then, they were stained with appropriate antibodies in FACS buffer (2% FBS in PBS) at 4°C for 30 minutes. The expression of the CAR was detected using a biotinylated protein L (GenScript, Piscataway, NJ) antibody and streptavidin-coupled PE (BD Biosciences, Franklin Lakes, NJ). The PE anti-human TGF-β Receptor II Antibody (clone W17055E, BioLegend, San Diego, CA) was used to detect dnTGFβRII expression on the CAR. The phosphorylation-SMAD2/3 ability of CAR T cells was detected using PE Mouse anti-Smad2/Smad3 antibodies (clone O72-670, BD Biosciences, Franklin Lakes, NJ) following the staining protocols. CD69-FITC (clone FN50, BioLegend, San Diego, CA) and CD25-PerCP/Cyanine5.5 (clone BC96, BioLegend, San Diego, CA) were used to detect T cell activation.

### Coculture Assay

In co-culture experiments, transduced or UTD T cells (2e5 cells per well in 100μL R10 media) were co-cultured with target cells (2e5 cells per well in 100μL R10 media) in 96-well round bottom tissue culture plates, at 37°C with 5% CO2 for 16 or 20hrs. In T cell phenotype assay, transduced or UTD T cells (5e5 cells per well in 500μL R10 media) were co-cultured with target cells (2.5e5 cells per well in 500μL R10 media) in 48-well flat bottom tissue culture plates. Target cells were irradiated with 10,000 rad ahead of co-culture with T cells. On day 2 and 4, 2.5e5 irradiated target cells were added in each well. After 5 days co-culture, remove 500ul supernatant from each well, gently resuspend and shift 250ul cells to 96well round bottom plate to stain. Human T cells were distinguished with live/dead viability stain (Thermo Fisher Scientific, Carlsbad, CA), followed by human CD3 and CD8 (clone OKT3 and clone SK1, BioLegend, San Diego, CA) stain. BV711-conjugated anti-human CD45RA (clone HI100, BioLegend, San Diego, CA), APC-conjugated anti-human CCR7 (clone G043H7, BioLegend, San Diego, CA) were used to detect T cell phenotypes. Staining was properly controlled with isotype antibodies. Before and after each staining, cells were washed twice with PBS containing 2% fetal bovine serum (FACS buffer). Fluorescence was assessed using a BD LSRFortessa flow cytometer and data were analyzed with FlowJo software.

### Mouse Model

All mouse experiments were conducted according to Institutional Animal Care and Use Committee (IACUC)-approved protocols. In orthotopic tumor model, 5e5 U87MG-CBG-GFP cells were implanted intracranially or 5e5 D270MG-CBG-GFP cells subcutaneously into 6- to 8- week-old NSG mice. The intracranial surgical implants were done using a stereotactic surgical setup with tumor cells implanted 2mm right and 2mm anterior to the lambda and 2mm into the brain. In subcutaneous models, NSG mice were injected with 5e5 D270 tumors subcutaneously in 100µL PBS on day 0. Tumor progression was evaluated by luminescence emission on a Xenogen IVIS spectrum after intraperitoneal injection of D-luciferin (Gold Biotechnology, St. Louis, MO) injection according to the manufacturer’s directions. Subcutaneous tumor was measured by calipers in length and width for the duration of the experiment. Tumor size was calculated as the area of tumor by multiplying the two dimensions. T cells were injected in a total volume of 100µL of PBS intravenously via the tail vein 7-8 days after tumor implantation. Survival was followed over time until a predetermined IACUC-approved endpoint was reached.

### Statistical Analysis

Data are presented as means ± SEM. RNA-seq data from TCGA was analyzed with Kruskal-Wallis test. Proliferation assay was analyzed with unpaired t test. Impedance and cytolysis assay were analyzed by ordinary one-way Analysis of Variance (ANOVA) with Tukey test to compare the differences in each group. The flow experiments were analyzed with two-way ANOVA with Tukey test to compare the differences in each group. Survival curves were analyzed with Kaplan-Meier (log-rank test). For the *in vivo* tumor study, linear regression was used to test for significant differences between the experimental groups. Survival, based on time to experimental endpoint, was plotted using a Kaplan-Meier curve. All statistical analyses were performed with Prism software version 9 (GraphPad, La Jolla, CA).

## ACKNOWLEDGEMENTS

The authors thank the Human Immunology Core at the University of Pennsylvania for providing T cells for the described work, the Stem Cell and Xenograft Core at the University of Pennsylvania for assistance with the animal work, the Small Animal Imaging Facility at the University of Pennsylvania for the bioluminescence imaging, the Daniel Martinez from Pathology Clinical Service Center of CHOP for immunohistochemistry staining, the Cell and Animal Radiation Core ((RRID SCR_022377), Flow cytometry core, the Histopathology Core, the BioRender to make the figure cartoon. Yunlin Zhang helped on the mice blood work. This work was supported by funding from GBM Translational Center of Excellence (N.L., D.M.O.), The Templeton Family Initiative in Neuro-Oncology (D.M.O.), The Maria and Gabriele Troiano Brain Cancer Immunotherapy Fund (D.M.O.), National Natural Science Foundation of China (U20A20383; Z.L.). N.L., J.L.R., Z.A.B., and D.M.O. are on patent filings related to the research presented here. DMO received monetary support from Tmunity Therapeutics for related lab work.

## AUTHOR CONTRIBUTIONS

D.M.O., Z.A.B., N.L. J.L.R., designed the experiments. N.L., J.L.R., Y.Y., M.T.L., L.Z., S.Y., K.A.H., Y.Z., L.Z., J.W., C.X., J.A.F., Z.A.B., Z.L., D.M.O. performed the experiments. N.L. and J.L.R., analyzed the data. N.L. wrote the manuscript with the help of ChatGPT to rewrite the grammar mistakes. D.M.O., Z.A.B., Z.L., provided funding. All the authors contributed to the editing and approval of the final manuscript.

