## Supplemental file for "Armored Bicistronic CAR T Cells with Dominant-negative TGF-β Receptor II to Overcome Resistance in Glioblastoma"

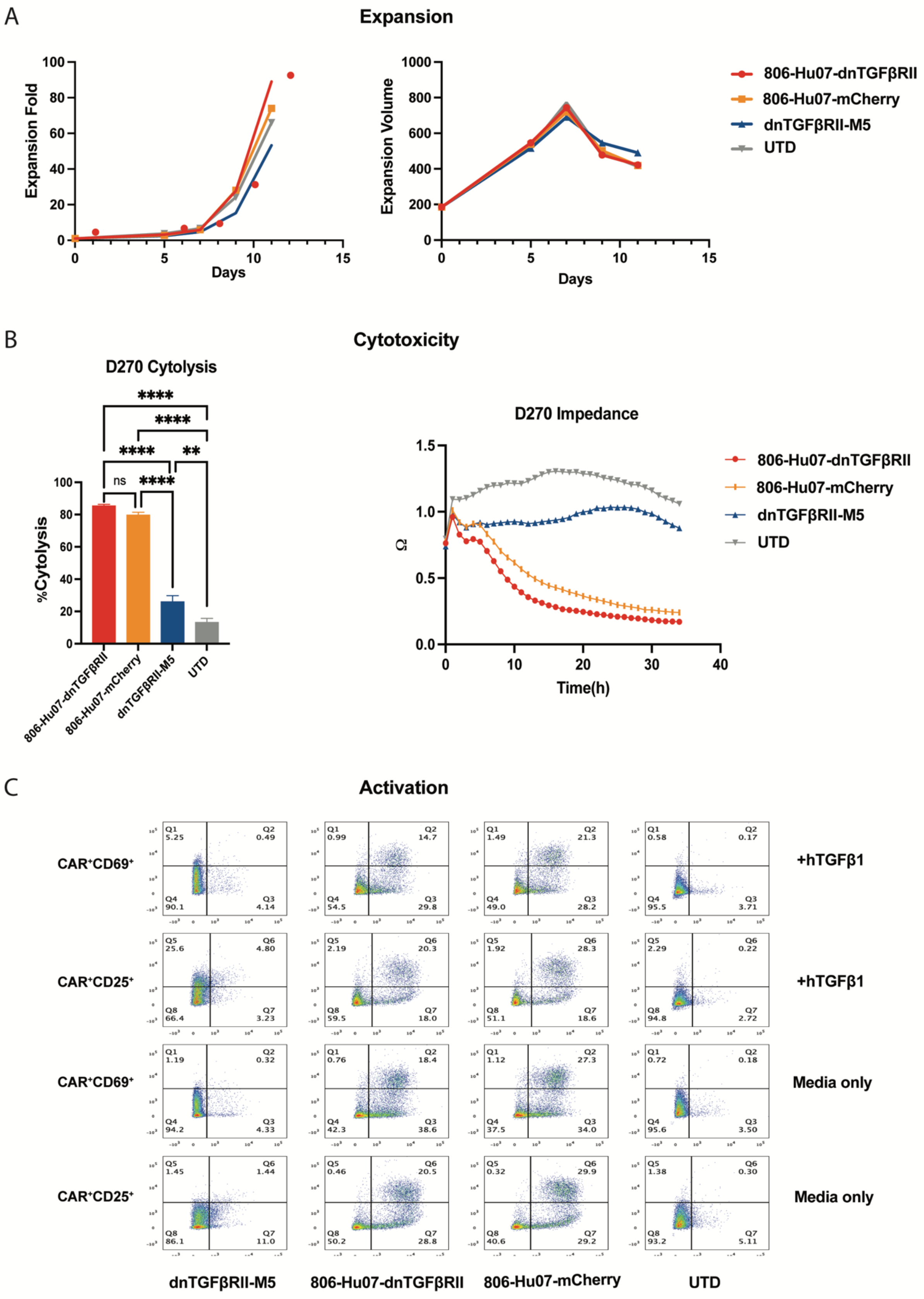
A
Expansion
Expansion Fold
Days
Expansion Volume
Days
806-Hu07-dnTGFβRII
806-Hu07-mCherry
dnTGFβRII-M5
UTD
B
Cytotoxicity
D270 Cytolysis
%Cytolysis
ns
****
**
806-Hu07-dnTGFβRII
806-Hu07-mCherry
dnTGFβRII-M5
UTD
D270 Impedance
Ω
Time(h)
806-Hu07-dnTGFβRII
806-Hu07-mCherry
dnTGFβRII-M5
UTD
C
Activation
CAR⁺CD69⁺
CAR⁺CD25⁺
CAR⁺CD69⁺
CAR⁺CD25⁺
+hTGFβ1
+hTGFβ1
Media only
Media only
dnTGFβRII-M5
806-Hu07-dnTGFβRII
806-Hu07-mCherry
UTD

**Figure S1. Kinetics of manufacturing T cells.**

(A) All the T cells exhibited a similar fold expansion and resting volume kinetics during the manufacturing. (B) Impedance cytotoxicity assays. Coculture D270MG cells with T cells, the data was collected every 1 min. (C) Flow cytometric detection of T cell activation markers. Scatter plot. Statistically significant differences were calculated by ordinary one-way ANOVA with Tukey's multiple comparisons test. ns, not significant; **p<0.01, ****p<0.0001. Data are presented as means ± SEM.

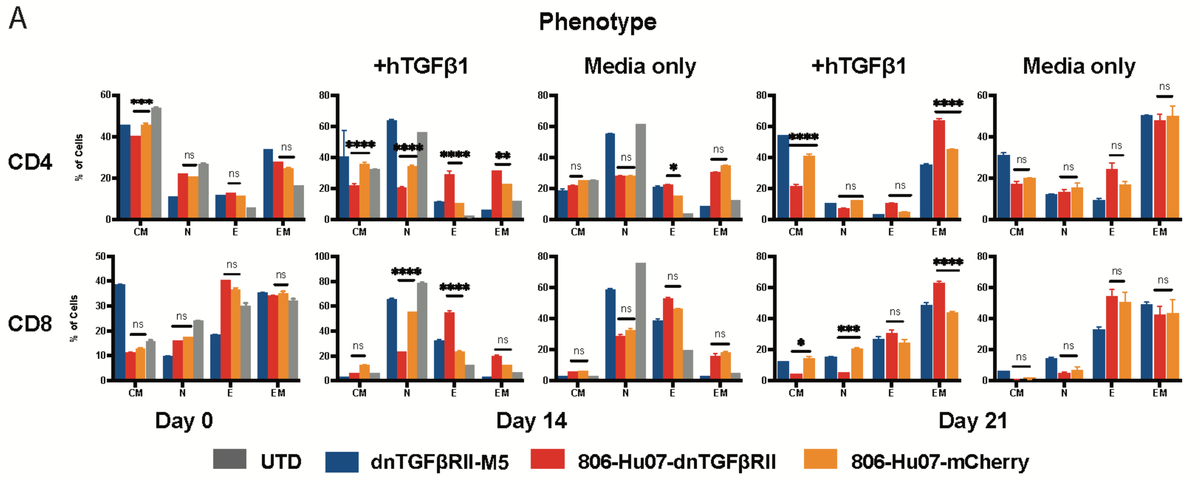


**Figure S2. T cells phenotypic evolution *in vitro* long-term restimulation assay.**

(A) They were collected on Day 0, Day 14, and Day 21 and stained with CD4 and CD8. The culture media was conditioned with or without hTGF-β1. These changes were evaluated within different subsets: Central Memory (CM), Naive (N), Effector (E), and Effector Memory (EM) T cells. Statistically significant differences were calculated by ordinary two-way ANOVA with Tukey's multiple comparisons test. ns denotes not significant, *p<0.05, **p<0.01, ***p<0.001, ****p<0.0001. Data are presented as the mean ± Standard Error of the Mean (SEM).
